# Irony detection engages the Precuneus and Inferior Frontal Gyrus and relies on integration of contextual cues and inferential skills

**DOI:** 10.1101/2022.08.15.504018

**Authors:** Elizabeth Valles-Capetillo, Cristian D. Ibarra, Magda Giordano

## Abstract

It has been suggested that irony is one of the most challenging forms of communication, consequently, it can be a valuable indicator of communication abilities. An ironic statement transmits the opposite meaning of its literal counterpart. The cognitive processes that may support the interpretation of irony include the Theory of Mind (ToM), executive functions, and processing style. The brain areas associated with irony detection are the medial prefrontal cortex, inferior frontal gyrus (IFG), posterior superior temporal gyrus (pSTG), precuneus, and inferior parietal lobule, among others. This study aims to analyze the cognitive processes and neural correlates involved in irony detection in Mexican adults. Forty-five participants underwent a cognitive assessment and performed a contextual discrepancy task during fMRI acquisition. The behavioral results showed that the detection of nonliteral statements (irony, unrelated, and white lies) requires ToM and verbal abilities. In addition, white lies detection seemed to involve inhibitory control. Ironic statements were the hardest intention to detect, having the lowest percentage of classification and the slowest latency of classification. Irony detection involved brain areas associated with the ToM (i.e., precuneus) and language (i.e., IFG and pSTG) as was expected based on the results of previous studies. The detection of literal and unrelated statements recruited motor areas. No differential activation pattern was found for detection of white lies. Finally, a global perceptual processing style predicted the percent change in the BOLD signal in the IFG for all the nonliteral and literal statements.

## Introduction

Language is used to communicate ideas, thoughts, and feelings through the combination of sounds, gestures, and symbols (1). Nevertheless, understanding the sounds that are part of a language, or the meaning of the words, is not enough to understand the richness of language’s meaning in social interactions (2). The ability to infer communicative intentions is a key process in human communication (3) because it allows the detection of the correct meaning between different possible interpretations of the same statement (4). It has been suggested that irony is one of the most challenging forms of communication (5) it can be a valuable indicator of communication abilities (6). According to the standard pragmatic view, an ironic statement can be defined as one that transmits the opposite meaning of its literal counterpart (7).

The cognitive processes that may support the interpretation of irony include the Theory of Mind (ToM), executive functions (8), and perceptual processing (2). The ToM, or mentalizing, refers to the inferences we make regarding mental states, a relevant ability since it is the mental states that determine actions (9). The relation between ToM and irony has been reported in behavioral studies in children with cerebral palsy (6). This relation has also been reported in neuroimaging studies, which have shown activation in areas associated with ToM (e.g., temporo parietal junction, medial prefrontal cortex, and the precuneus) during irony detection (10). Executive functions (EF) are general-purpose control mechanisms regulating human cognition and action (11). There are three core EFs, inhibition, working memory, and cognitive flexibility, from which higher-order EFs are built (12). Inhibition refers to inhibitory control, including self-control (behavioral inhibition) and interference control (selective attention and cognitive inhibition) (12). The three core EFs have been associated with the detection of irony in older adults (13). In typically developing children, inhibitory control has been associated with irony comprehension (6). Additionally, pragmatic language requires the use of context to derive meaning; thus, the ability to integrate the utterance with its surrounding context is crucial (2). The ability to integrate sources of information has been termed “central coherence,” weak central coherence (WCC) results in attention being given to small pieces at the expense of coherent global patterns of information (2). WCC is a bias in a cognitive style that is evident from the perceptual level (visual illusions) to the most complex tasks, such as pragmatic understanding (2,14).

Pragmatic language difficulties can be found in various clinical populations, including traumatic brain injury, right-hemisphere damage, schizophrenia, neurodegenerative disorders, attention-deficit hyperactivity disorder (ADHD), and Autism Spectrum Disorder (ASD) (2,6,14,15). Studies in these clinical populations help clarify the role that ToM, EFs, intellectual capacity, and cognitive biases play in pragmatic language abilities. It should be noted that among non-clinical individuals, there is variability in those aspects that appear to be affected in the clinical population. One example is central coherence, which varies from strong—with a focus on meaning or gestalt—to weak—with a focus on details—in the general population (16). In their study, Boot and Happé (2010) evaluated whether differences in detail focus could be separated from differences in intellectual ability in typically developing children and adolescents (8-25 years of age; mean age 14.5 years). To measure central coherence, they used a verbal task that required participants to complete sentence stems. In this sample, they found that a significant proportion of the variance in completion scores could not be explained by age, intellectual quotient, or gender. This result could reflect individual differences in cognitive style. Another example is social-communication disability which has been proposed to vary between autism and normality (17). Baron-Cohen et al., (2001) designed a brief, self-administered questionnaire, the *Autism Spectrum Quotient (AQ),* to evaluate “where any given individual adult, with normal intelligence, lies on this continuum”. The AQ evaluates five different areas, i.e. social skills, communication, attention to detail, attention switching, and imagination. Baron-Cohen et al., (2001) found that participants with a diagnosis of Asperger/High Functioning Autism scored significantly higher than matched controls, that within the control group, men scored higher than women, and that there were differences among occupations, replicating earlier findings from the group. Specifically, the authors found that scientists scored higher than nonscientists; and that “within the sciences, mathematics, physical scientists, computer scientists, and engineers scored higher than the more human or life-centered sciences of medicine (including veterinary science) and biology”.

The brain areas that have been associated with irony detection are the medial prefrontal cortex (MPFC)—either bilaterally or unilaterally—(10,18–22). In the left hemisphere, the inferior frontal gyrus (IFG) (10), the inferior parietal lobe (IPL) (10,23), pre-supplementary and motor area (18), precentral gyrus (23), and precuneus (18). And in the right hemisphere, the precentral gyrus (18,23), the dorsolateral prefrontal cortex (dlPFC) (10), the IFG (10), temporal superior gyrus (22) and the cingulate gyrus (19,23). The results described above come from articles with significant methodological variability concerning the native language of the participants, the modality of the stimuli, and the writing system; none included Spanish-speaking participants. Despite the methodological variability, a meta-analysis showed that the dorsomedial prefrontal cortex (dmPFC), rostromedial prefrontal cortex (rmPFC), IFG, and pSTG presented consistent activation during irony detection (24).

The present study aimed to analyze the cognitive processes involved in irony detection, including social cognition (e.g., ToM), EF, perceptual processing, and verbal abilities, and find the neural correlates in a sample of neurotypical Mexican adults. To increase the variability in the sample, participants from different professional categories (i.e., science, humanities, and engineering) were included. We expected to replicate and expand the findings of the meta-analysis (24) in terms of the brain regions involved in irony detection and to improve on the previous studies by including a thorough cognitive evaluation of the participants.

## Materials and Methods

### Participants

Participants were asked to fill out a general data form with information about their level of education, sex, and age. The experiment was completed by 45 participants (22 female) with a mean age of 26.69 ± 5.83. Of the 45 participants, 15 were studying or had a degree related to science (7 female), 12 non-scientific disciplines (e.g. humanities, 6 female), and 18 engineering (9 female). Participants reported no history of psychiatric or neurological illness, were right-handed, and signed a written informed consent to participate in the study. Handedness was measured with the Edinburgh handedness inventory (25). The experimental protocol was approved by the Ethical Committee of the Neurobiology Institute of the Universidad Nacional Autónoma de México (#047.H.RM).

### Behavioral paradigm

To assess irony detection, a contextual discrepancy task for Mexican adults was created (26). Fifty-six social contexts and 14 statements were constructed. Each social context category—i.e., ironic, white lies, literal and unrelated–––generated an environment that modified the intention of the statement that followed. Each statement was used with all four categories of social contexts. The task was created in Psychopy (3.0.1 version) (27), and the stimuli were presented in the auditory modality without prosodic changes (see Fig 1). The participants heard the social context, then the statement, and in the last screen, a question regarding the classification of the statement was presented in text modality, with four options.

**Figure 1.-.**
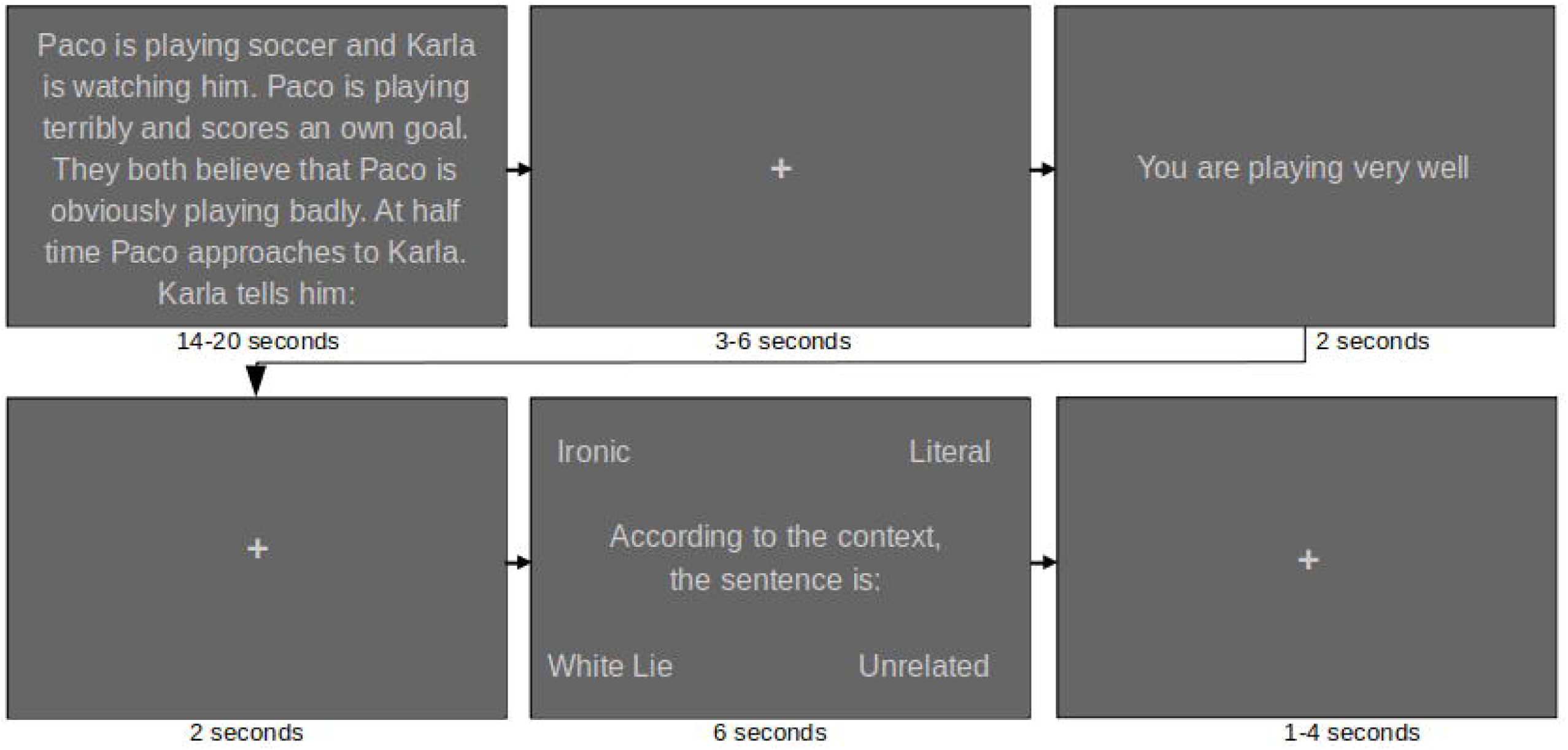
Contextual discrepancy task. The context and statement were presented in audio without changes in prosody (note: text was not presented). The classification screen was presented in text modality. Stimuli were presented in semi-random order (26).

### Cognitive tests

A battery of tests was used to evaluate the cognitive processes that could be associated with irony detection. The cognitive variables of interest were social cognition, executive functions, perceptual processing, and verbal abilities. For social cognition tests, we used tests that evaluate ToM, empathy, use of sarcasm, and social communication abilities. ToM was assessed with the Short Story Task (SST) (28) and the Reading the Mind in the Eyes Test (RMET) (29). Empathy was assessed with the Interpersonal Reactivity Index (IRI), which has four scales: fantasy, perspective-taking, empathic concern, and personal distress (30). To assess the degree to which an individual is likely to use sarcasm, the sarcasm self-report scale was used (SSS) (31). The AQ was selected to assess the degree of social abilities, communication, attention to detail, attention switching, and imagination; this inventory has proved to be sensitive in non-clinical populations as described before (29).

For executive functions, the Psychology Experiment Building Language software (PEBL) was used (32). The EF evaluated were: updating, which was assessed with the n-back test; planning, assessed with the Tower of London; inhibitory control, was assessed with the go-no-go test; and for working memory, digit span test. Perceptual processing was assessed with the Perceptual Reasoning Index (PRI) of the Wechsler Adult Intelligence Scale, fourth edition (WAIS-IV) (33) and the local-global test (34) using Psychtoolbox-3 (35). The local-global task has been used to evaluate WCC with neurodevelopmental disorders, such as ASD and Williams syndrome (36). Verbal abilities were assessed with the verbal comprehension index (VCI) from the WAIS-VI (33); and verbal fluency, assessed with the BANFE (Batería Neuropsicológica de Funciones Ejecutivas) (37).

A Shapiro test showed that the results of the behavioral task were not normally distributed (p < 0.01). Levene’s test showed that the variances between the statement categories were unequal (p < 0.01). Therefore, non-parametric statistics were used. The percentages and standard deviation are presented for the classification accuracy, and the median and interquartile range (IQR) for the latency of classification. A comparison between statement categories was performed using a Friedman test for repeated measures and a Durbin-Conover pairwise comparison analysis with Bonferroni correction.

To find out which were the best cognitive predictors of behavioral performance during the contextual discrepancy task, we used a stepwise regression model for accuracy and latency of classification using R. The stepwise regression starts with no predictors, then sequentially adds the most contributive predictors, then removes the variables that do not improve the model fit (38). The predictors were selected by category of the cognitive process evaluated: social cognition, executive functions, perceptual processing, and verbal abilities. For social cognition, we included the total score for the RMET, SST, IRI, SSS, and AQ. In addition, we included reaction time (rt) for the RMET. For EF, we included the go-no-go, Tower of London, digit span, and n-back. For perceptual processing, we included the local global mean accuracy for switching and non-switching and the difference in the rt between switching and non-switching, and the PRI. And for verbal abilities, verbal fluency and VCI were selected.

### Functional Magnetic Resonance Imaging

The MRI session was performed on a 3-Tesla General Electric MR750 scanner with a 32-channel head coil. The acquisition parameters were, for T1w: TE of 0.003 seconds, TR of 0.008, flip angle of 12°, and a 256 x 256 matrix; for the functional sequence: TE of 0.04, TR of 2 seconds, 90° flip angle, and 64 x 64 matrix.

The preprocessing from dicom to nifti format conversion was performed using dcm2niix v1.0.20190902 (39). The preprocessing of the MRI data was performed using the fmriprep pipeline (40).

The contextual discrepancy task presentation was optimized for the MRI scanner session using OPTSEQ (41) and divided into three runs. Each run had a duration of 10 minutes. We asked the participants to infer the speaker’s intention when they heard the statement and to use the information provided by the context to do this task. For the participants to become familiar with the task, before starting the task within the scanner, a training task was given in which they classified 4 stimuli (one per category). The stimuli of the training task were different from those used in the task inside the scanner. NordicNeuroLab goggles (Bergen, Norway) were used for task presentation and a response device (Cedrus Lumina, California, USA), both compatible with the scanner.

A whole brain analysis, for the subsequent analysis, was performed with the FMRI Software Library (FSL; v. 6.0.5) (42). For the first (run), second (participant), and third (group) level analyses, a general linear model (GLM) analysis was performed using the FEAT tool (43). In the first level analysis, the BOLD signal was analyzed during the presentation of the event of interest (i.e., statements). To analyze the events, a GLM was performed using 15 regressors, four for each statement category (i.e. irony, literal, unrelated, and white lies), one for fixation cross, and 11 for physiological regressors. The physiological regressors were cerebrospinal fluid, white matter, global signal, dvars, frame wise displacement, and 6 motion regressors (translation and rotation of the x, y, z coordinates). Because fmriprep does not perform smoothing, a 6 mm smoothing was applied at this level, and the significance threshold was adjusted at the cluster level (z = 2.3 and p < 0.05). To analyze the differences between the statement categories, 11 contrasts were analyzed: 4 contrasts comparing each statement category with the fixation cross (e.g. irony > fixation cross), 4 comparing each statement category with the other three (e.g. irony > white lies + literal + no relation), and 3 comparing irony with other statement categories (e.g. irony > white lies).

For the second level analysis (subject level), a fixed effects GLM was performed. And for the third-level analysis (group level), a mixed-effects GLM (i.e. Flame 1) was performed. For both analyses, the threshold parameters of the first level analysis were maintained with cluster correction (z = 2.3, p < 0.05). The functional activation areas (i.e. labels) were identified using the corresponding MNI152 template coordinates. The labels were obtained using the MNI <-> Talairach tool with the Brodmann areas in the Bioimage Suite Web (44). This tool defined the Brodmann areas in MNI space (45).

To analyze the relationship between the activation in the brain during comprehension of the statements and the cognitive processes we measured, the percentage of BOLD signal was extracted, per subject and condition, from regions previously associated with irony using the feat_query tool. To obtain the percent of BOLD signal we selected the contrast that compared each statement category (ironic, white lies, literal and unrelated) with the fixation cross (e.g. ironic statements > fixation cross). The Regions of Interest (ROIs) were selected from a meta-analysis that studied irony (24); they were the dmPFC (x= 2, y = 54, z= 20), IFG (x= −50, y = 16, z= 4), pSTG (x= −60, y = −40, z= 12), and rmPFC (x= 2, y = 56, z= 16). The ROIs had a radius of 6 mm and were constructed using the fslmaths function.

To find out which were the best cognitive predictors of the BOLD signal during the presentation of each statement category, we used a stepwise regression model using R. The predictors were selected by category of the cognitive process evaluated: social cognition, executive functions, perceptual processing, verbal abilities, as described with regard to the behavioral performance in the contextual discrepancy task.

## Results

### Behavioral

#### Contextual discrepancy task

The ironic statements showed the lowest percentage of correct answers (71.43 ± 28.16), followed by white lies (78.57 ± 20.04), and literal (85.71 ± 13.52), while unrelated statements showed the highest accuracy (92.86 ± 14.64). Ironic statements required the longest latency of classification (seconds) (Md = 1.91, IQR = 1.01), followed by literal (Md = 1.72, IQR = 0.87), white lies (Md = 1.49, IQR = 1.20), and unrelated statements (Md = 1.31, IQR = 0.84).

The results show that there are significant differences in the percentage of correct answers (*X*^2^_Friedman_(3) = 45.68, p < 0.001) and latency of classification (*X*^2^_Friedman_(3) = 31.81, p < 0.001). In the percentage of correct answers, significant differences were found between the statement categories of irony compared with literal (p < 0.05), unrelated (p < 0.001), and white lies (p < 0.001); also, between the unrelated statements compared with literal (p < 0.001) and white lies (p < 0.001; see Fig 2A). Latency of classification showed significant differences between the ironic statement compared to unrelated (p < 0.001) and white lies (p < 0.001), and also between literal compared with unrelated (p < 0.001) and white lies (p < 0.001; see Fig 2B).

**Figure 2.-.**
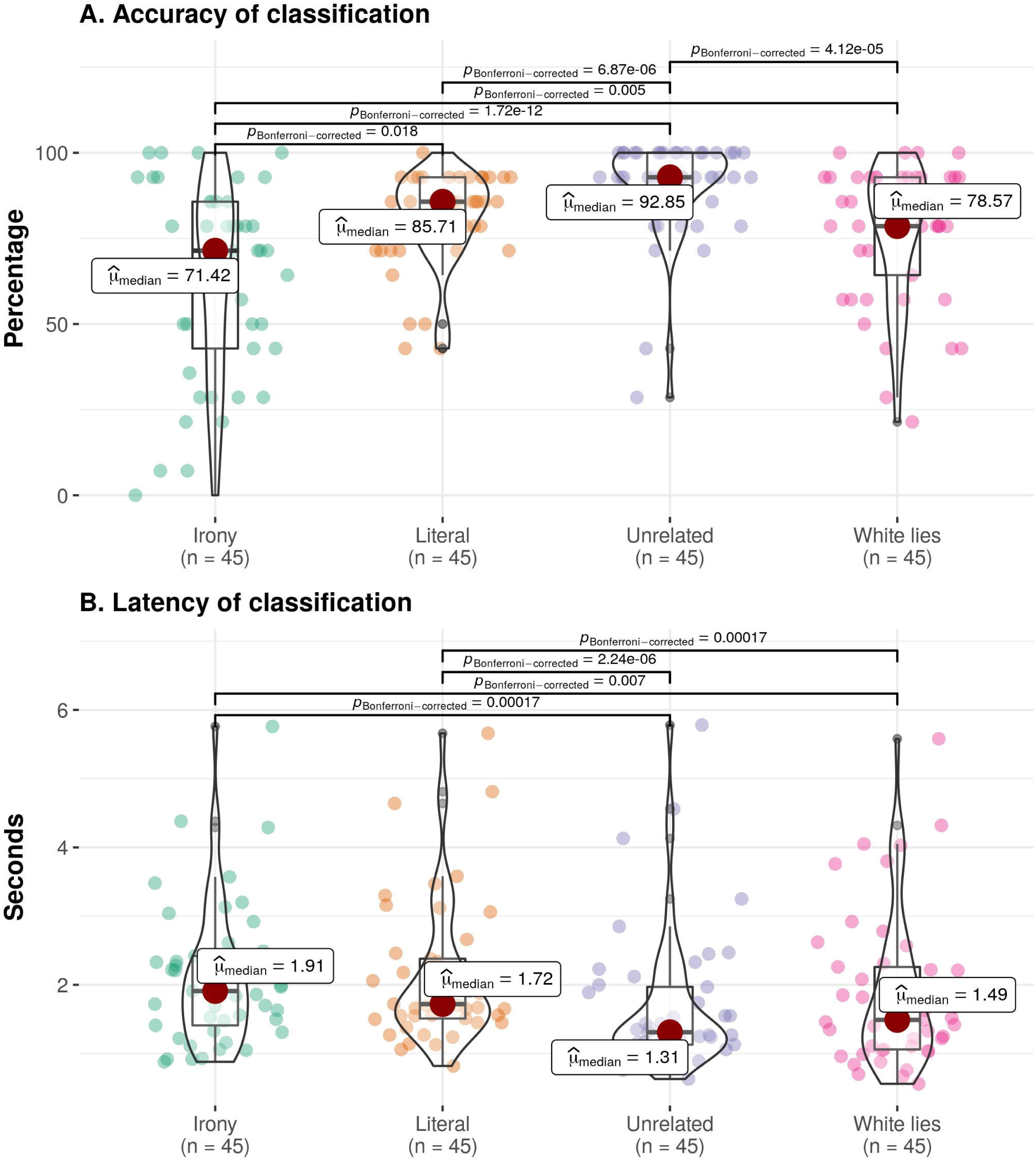
Percentage of correct answers and latency of classification for each statement category. **A**. Ironic statements resulted in a significantly lower percentage of correct answers, followed by white lies, and literal, while unrelated, had the highest percentage of correct answers. **B**. Latency of classification showed significant differences between the statement categories of irony compared with unrelated and white lies, and also between literal compared with unrelated and white lies, The plots show the density curves, and the box plots show the median (red circle), mean (thick line), interquartile range (rectangle), and the lower/upper adjacent values (black lines stretched from the rectangle), and scatter plot. The significant differences between statement categories are reported with the p-value.

#### Contextual discrepancy task and cognitive processes

As described in the methods, we used a stepwise regression model for accuracy and latency of classification to find out which were the best cognitive predictors of behavioral performance during the contextual discrepancy task. The predictors were selected by category of the cognitive process evaluated: social cognition, executive functions, verbal abilities, and perceptual processing (Table 1, and S1 Table).

**Table 1.** Stepwise regressors for behavioral results, by cognitive processes and statement categories.

| Model./<br>dependent.varia<br>ble | Social.cog<br>nition | Executive.fu<br>nctions | Perceptual.<br>processing | Verbal.abilit<br>ies |
| --- | --- | --- | --- | --- |
| Irony |  |  |  |  |
| Accuracy | 0.12* | 0.03 | n.s. | 0.14** |
| Latency.of.classif<br>ication | 0.21** | 0.04 | 0.03 | n.s. |
| Literal |  |  |  |  |
| Accuracy | n.s. | n.s. | 0.02 | n.s. |
| Latency.of.classif<br>ication | 0.083 | n.s. | 0.02 | 0.027 |
| Unrelated |  |  |  |  |
| Accuracy | 0.34*** | n.s. | n.s. | 0.19*** |
| Latency.of.classif<br>ication | 0.09* | n.s. | n.s. | n.s. |
| White.lies |  |  |  |  |
| Accuracy | 0.078* | 0.067* | n.s. | 0.20*** |
| Latency.of.classif<br>ication | 0.16** | 0.07* | 0.02 | n.s. |

The adjusted-R<sup>2</sup> are presented for each model. For details on the analysis see Supplementary Table 2.
p = \* < 0.05, \*\* < 0.01, \*\*\* < 0.001, n.s. = no significant predictors survived in the stepwise regression

Accuracy of irony detection was predicted by scores in AQ (t = −1.55, p = 0.13), and RMET (t = 2.588, p < 0.05); in combination, they made a significant contribution (F(2,41) = 4.16, p < 0.05, adjusted-R2 = 0.12). Also by VCI (t= 2.83, p < 0.01; F(1,41) = 8.01, p < 0.01, adjusted-R2 = 0.14). Similarly, the accuracy of unrelated statements detection was predicted by scores in AQ (t = −2.37, p < 0.05) and RMET (t = 4.54, p < 0.0001); in combination, they made a significant contribution (F(2,41) = 12.1, p < 0.0001, adjusted-R2 = 0.34). VCI (t= 3.41, p < 0.01) also predicted the accuracy of unrelated statements detection (F(1,41) = 3.42, p < 0.01, adjusted-R2 = 0.19). Accuracy of white lies detection was predicted by scores in RMET (t = 1.27, p < 0.05; F(1,42) = 4.64, p < 0.05, adjusted-R2 = 0.08), scores in the go-no-go task (t = 2.02, p < 0.05) (F(1,42) = 4.10, p < 0.05, adjusted-R2 = 0.07), and VCI (t= 3.45, p < 0.01; F(1,41) = 11.93, p < 0.01, adjusted-R2 = 0.20). For accuracy of literal statements detection, there were no significant predictors (Table 1, Figs 3A and 3B, and S2 Table).

**Figure 3.**
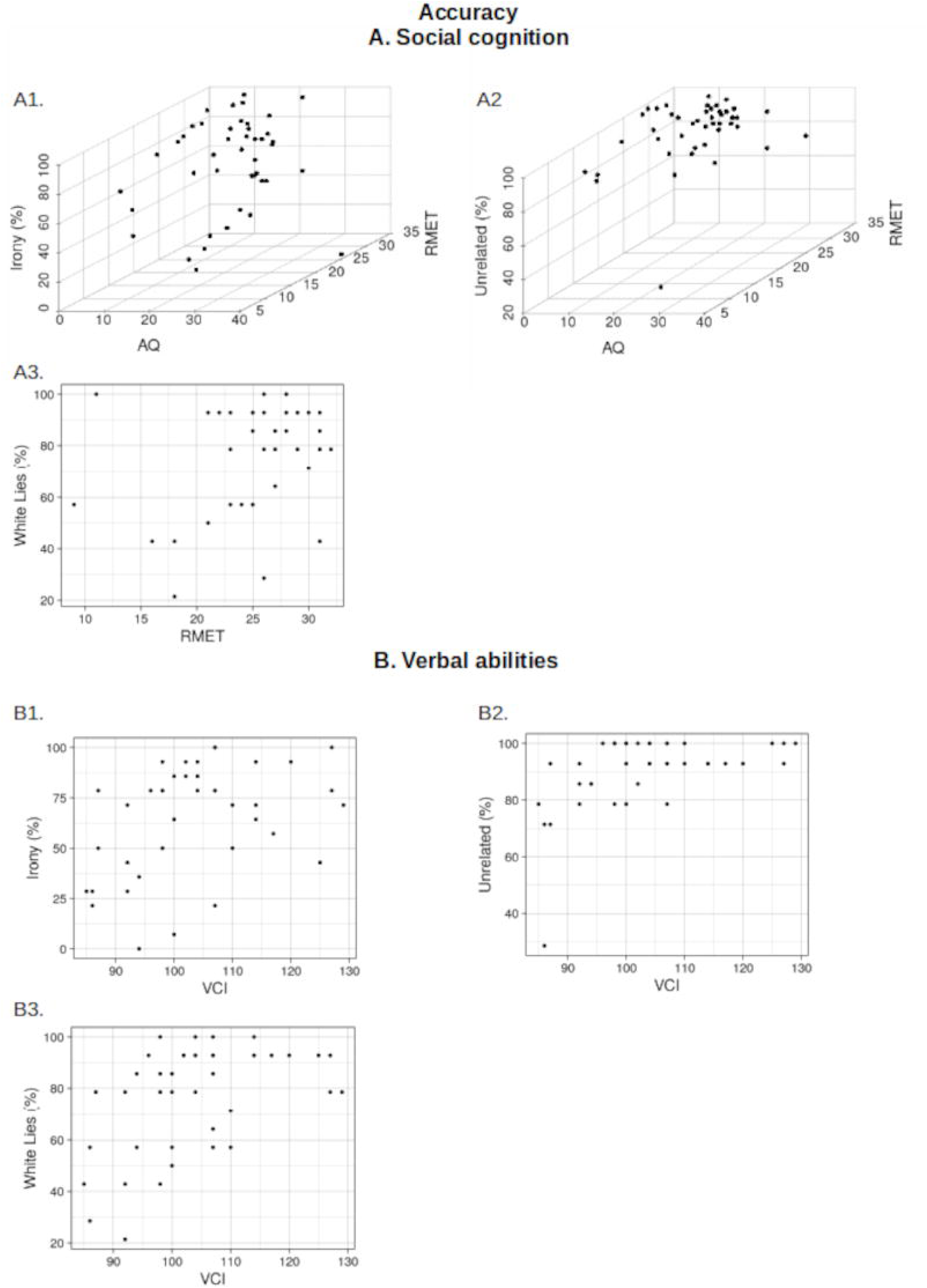
Figures show the relationship between the cognitive predictors’ scores and the behavioral results of the contextual discrepancy task (percent accuracy). Panel A shows the role of social cognitive processes (AQ and RMET). Panel B showed the role of verbal abilities, i.e., VCI. For statistical results see: section 3.1.2. AQ: autism quotient; RMET: reading the mind in the eyes task, VCI: verbal comprehension index from the WAIS-IV.

For latency of classification, scores in AQ (t = 3.27, p < 0.01) and SST (t = 3.30, p < 0.01) predicted latency of classification of ironic statements; in combination they made a significant contribution (F(2,41) = 6.83, p < 0.001, adjusted-R2 = 0.21). Scores in AQ (t = 2.19, p < 0.05) and SST (t = 1.61, p = 0.11) predicted latency of classification of unrelated statements; in combination they made a significant contribution (F(2,41) = 3.13, p < 0.05, adjusted-R2 = 0.09). Similarly, scores in AQ (t = 2.82, p < 0.01) and SST (t = 2.16, p < 0.05) predicted latency of classification of white lies; in combination they made a significant contribution. Scores in the go-no-go task (t = −2.09, p < 0.05) also predicted latency of classification of white lies (F(1,41) = 4.37, p < 0.05, adjusted-R2 = 0.07). For literal statements, there were no significant predictors (see Table 1 and S2 Table).

### Functional Magnetic Reasoning Imaging

#### The contrast between one statement category versus the others

The contrast irony > literal+unrelated+white lies showed significant differential activation in the precuneus, in Brodmann’s area 31 (BA) (see Fig 4A; Table 2). The contrast literal > irony+unrelated+white lies showed significant differential activation in three clusters: the right premotor area (PrM; BA 6); the left primary motor area (PMA; BA 4), extending to the primary sensory area (BA 1); and the right putamen, extending to the caudate (see Fig 4B; Table 2). The contrast unrelated > irony+literal+white lies showed significant differential activation in three clusters: the left premotor area (BA 6), extending to the anterior cingulate gyrus (BA 24); the left visual-motor area (BA 7), extending to the anterior supramarginal gyrus (BA 40); and the right primary motor area (BA 4), extending to the premotor area (BA 6) (see Fig 4C; Table 2). The contrast white lies > irony+literal+unrelated did not show significant results.

**Figure 4.-.**
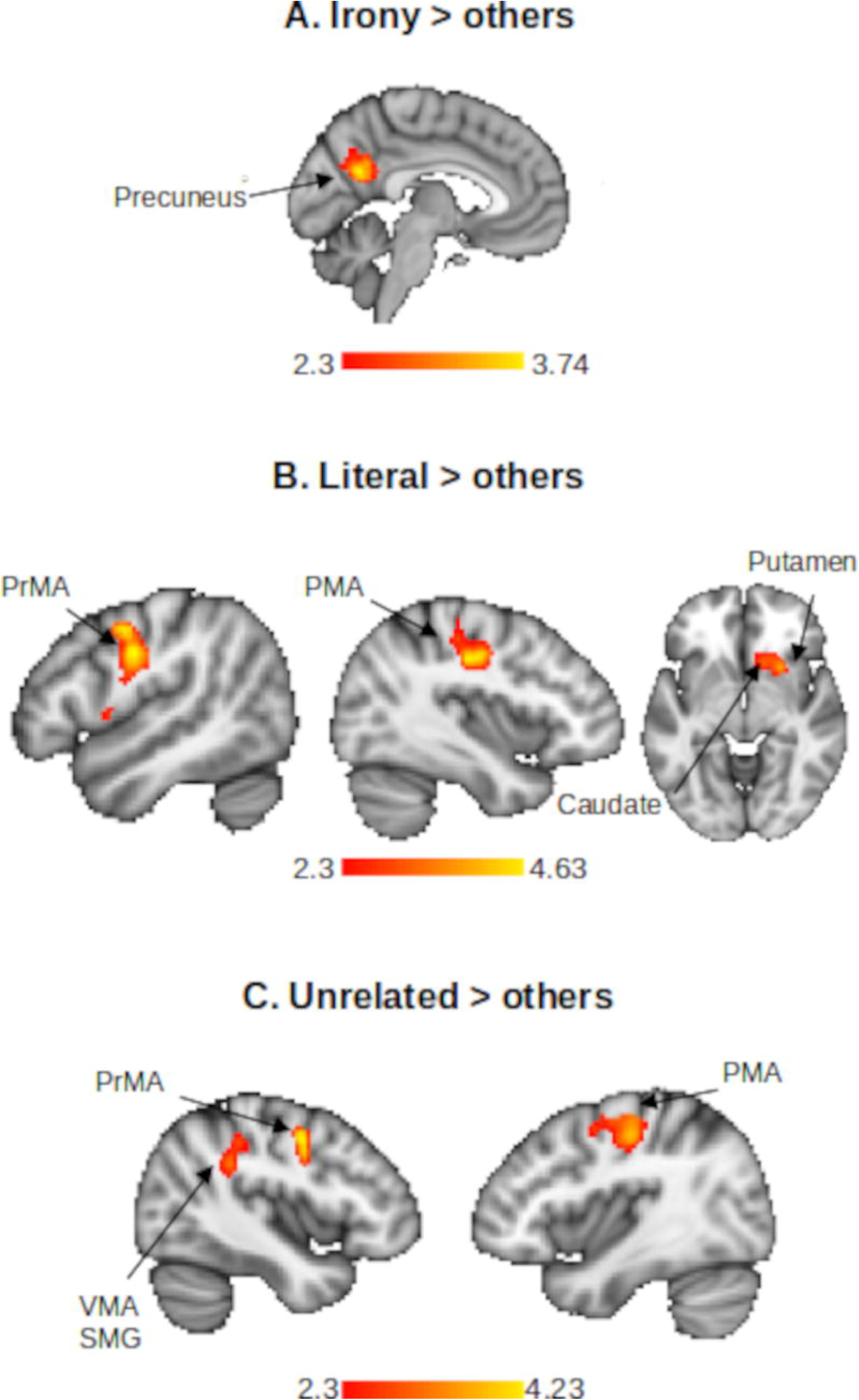
Results of the contrast between each statement category and the others. Panel A shows the contrast irony > literal+unrelated+white lies, this contrast shows significant differential activation in the precuneus. Panel B shows the contrast literal > irony+unrelated+white lies. Panel C shows the contrast of unrelated > irony+unrelated+white lies. The contrast white lies > irony+literal+unrelated did not show significant differential activation. Maps were thresholded at z < 2.3, p < 0.05. PrMA = premotor area, PMA = primary motor area, VMA = ventral motor area, SMG = supramarginal gyrus.

**Table 2.**
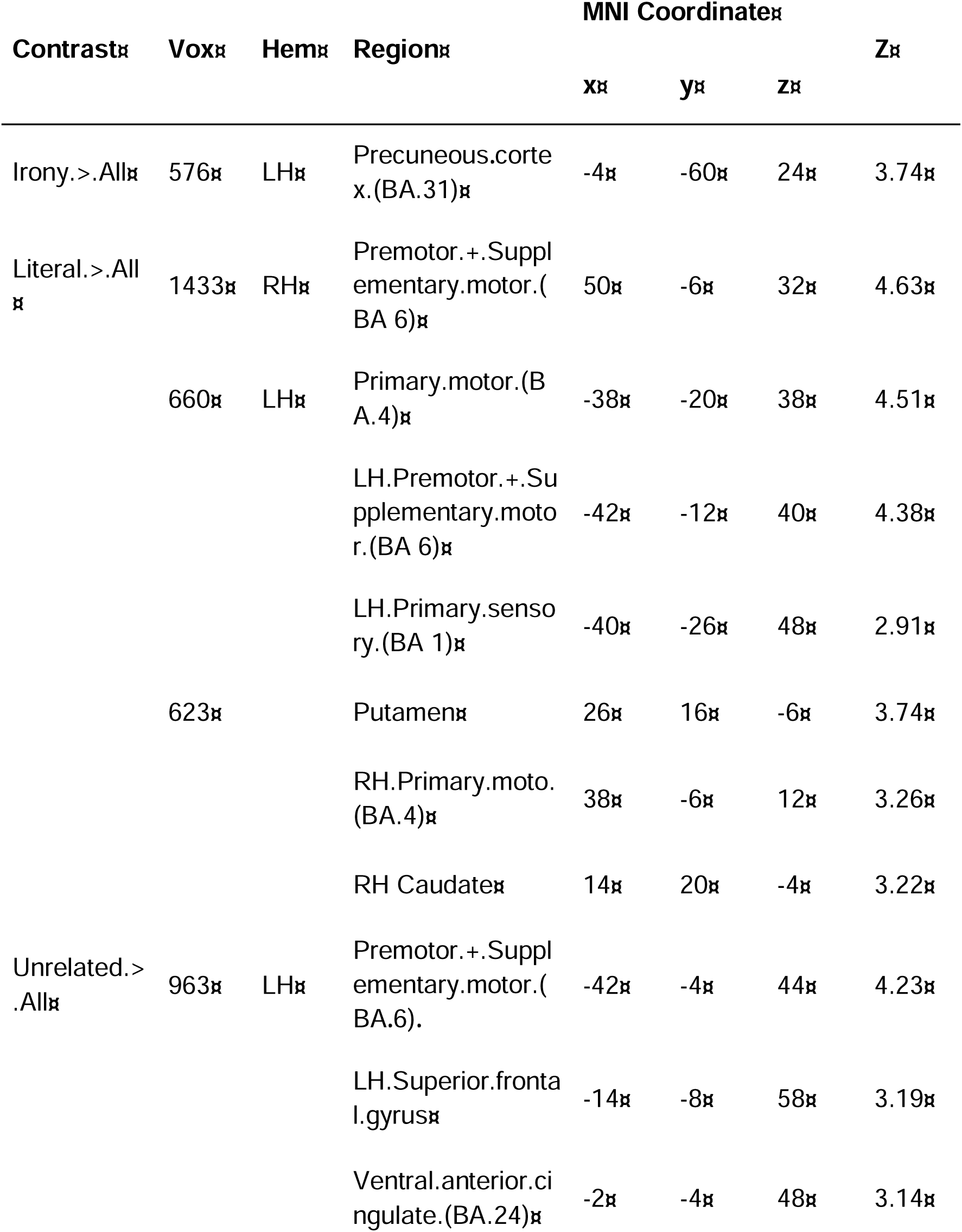
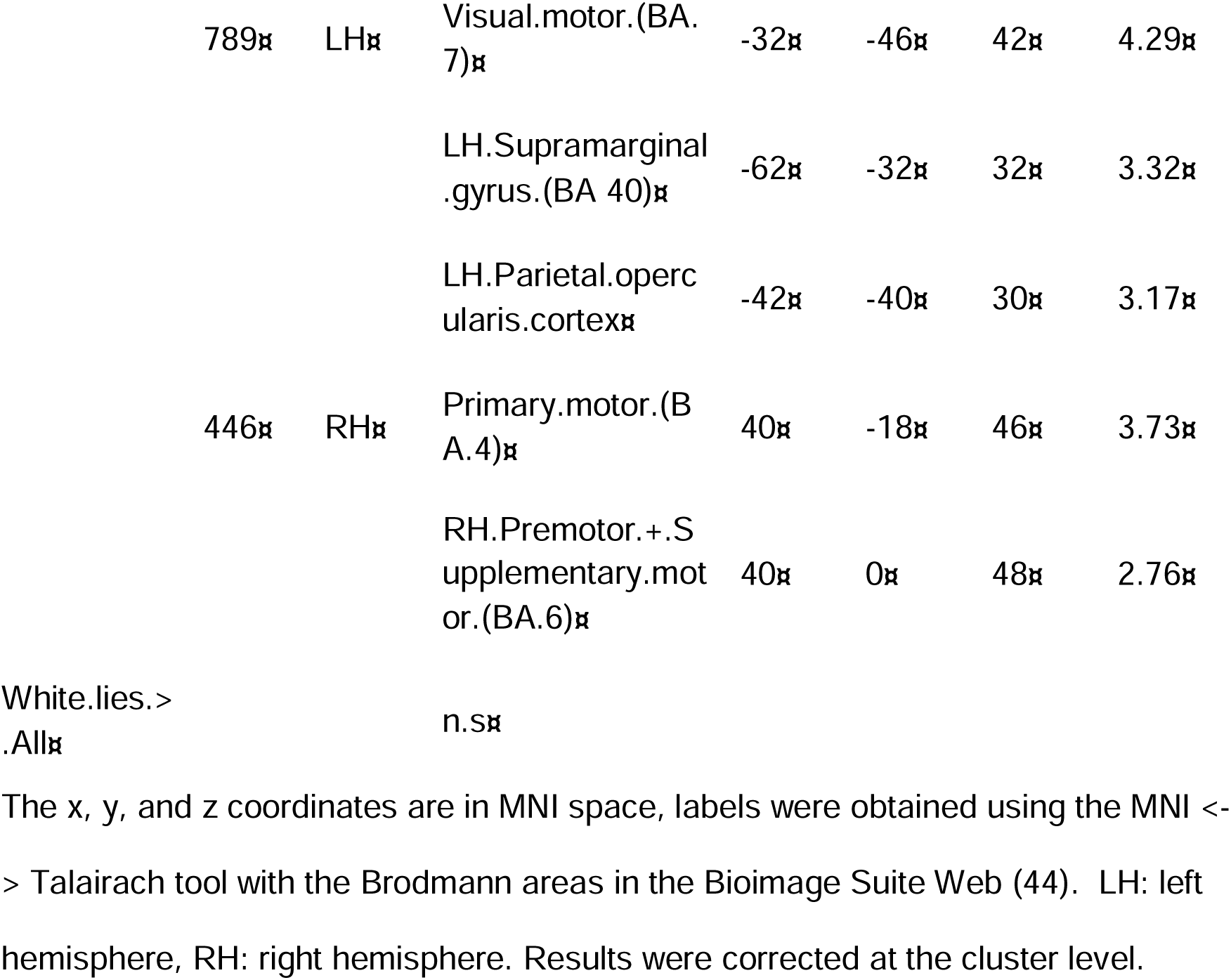
Brain areas exhibiting significant activation in whole brain analysis for contrasts between one statement category versus the others, according to GLM analysis. The x, y, and z coordinates are in MNI space, labels were obtained using the MNI <-> Talairach tool with the Brodmann areas in the Bioimage Suite Web (44). LH: left hemisphere, RH: right hemisphere. Results were corrected at the cluster level.

#### The contrast between ironic statements and other statement categories

The irony > literal contrast showed significant differential activation in three clusters, the precuneus, extending into the left motor visual area (BA 7) and the right posterior superior temporal gyrus visual-motor area (BA 7); the left angular gyrus (BA 39), extending into the medial temporal gyrus (BA 31) and pSTG (BA 22); and the right IFG (pars triangularis, BA 45,), extending into the dorsolateral prefrontal cortex (BA 9) and IFG (pars opercularis, BA 44) (see Fig 5A; Table 3 for MNI coordinates).

**Figure 5.-.**
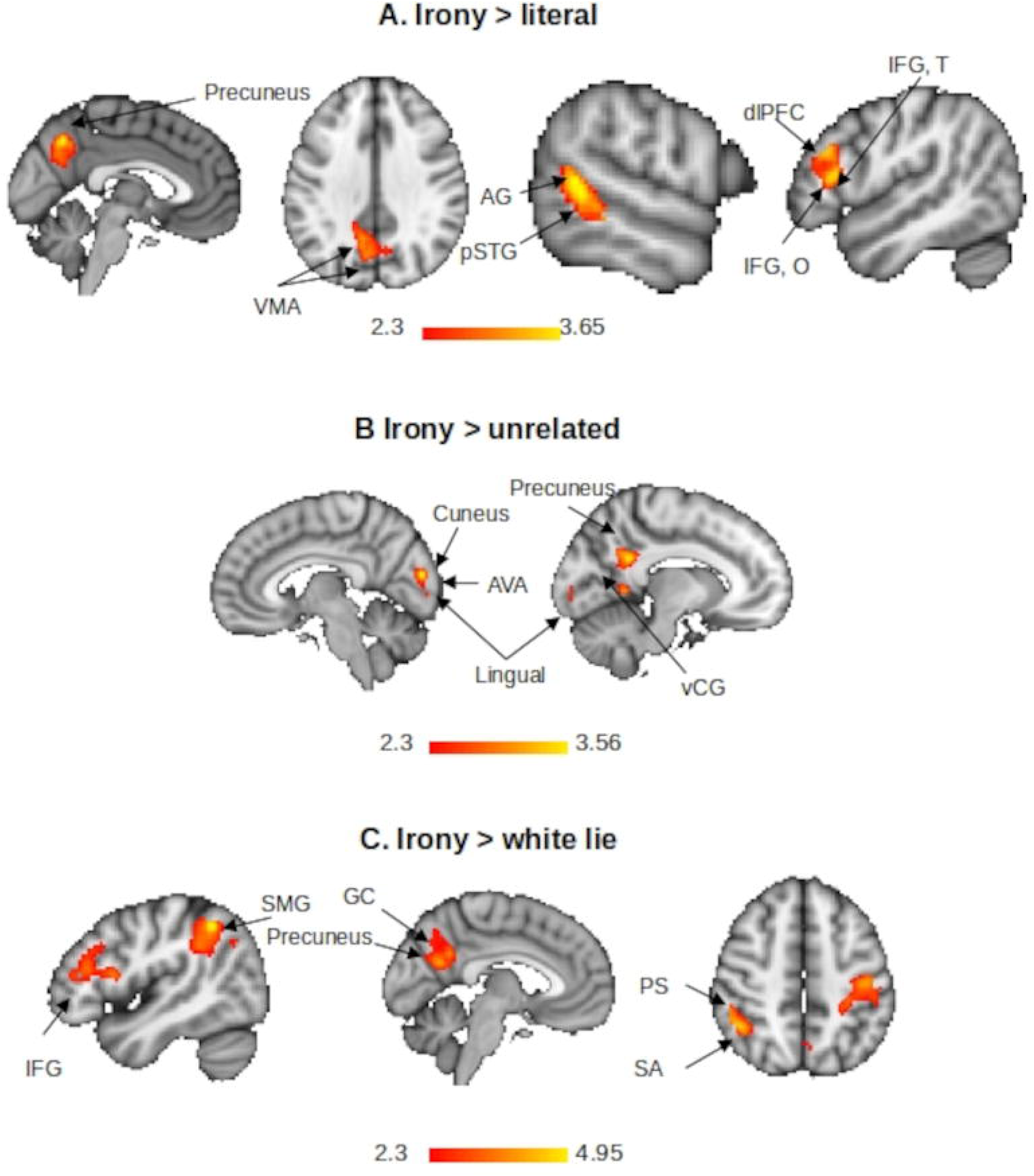
Contrasts of irony versus other statement categories. Panel A shows the contrast irony > literal; Panel B, irony > unrelated; panel C, irony > white lies. Maps were thresholded at z < 2.3, p < 0.05. VMA = visual motor area, AG = angular gyrus, pSTG = posterior superior temporal gyrus, dlPFC = dorsolateral prefrontal cortex, IFG T = inferior frontal gyrus triangularis, IFG O = inferior frontal gyrus opercularis, AVA = associative visual area, VCG = ventral cingulate gyrus, SMG = supramarginal gyrus, GC = cingulate gyrus, SA = primary association cortex, PS = primary sensory cortex.

**Table 3.**
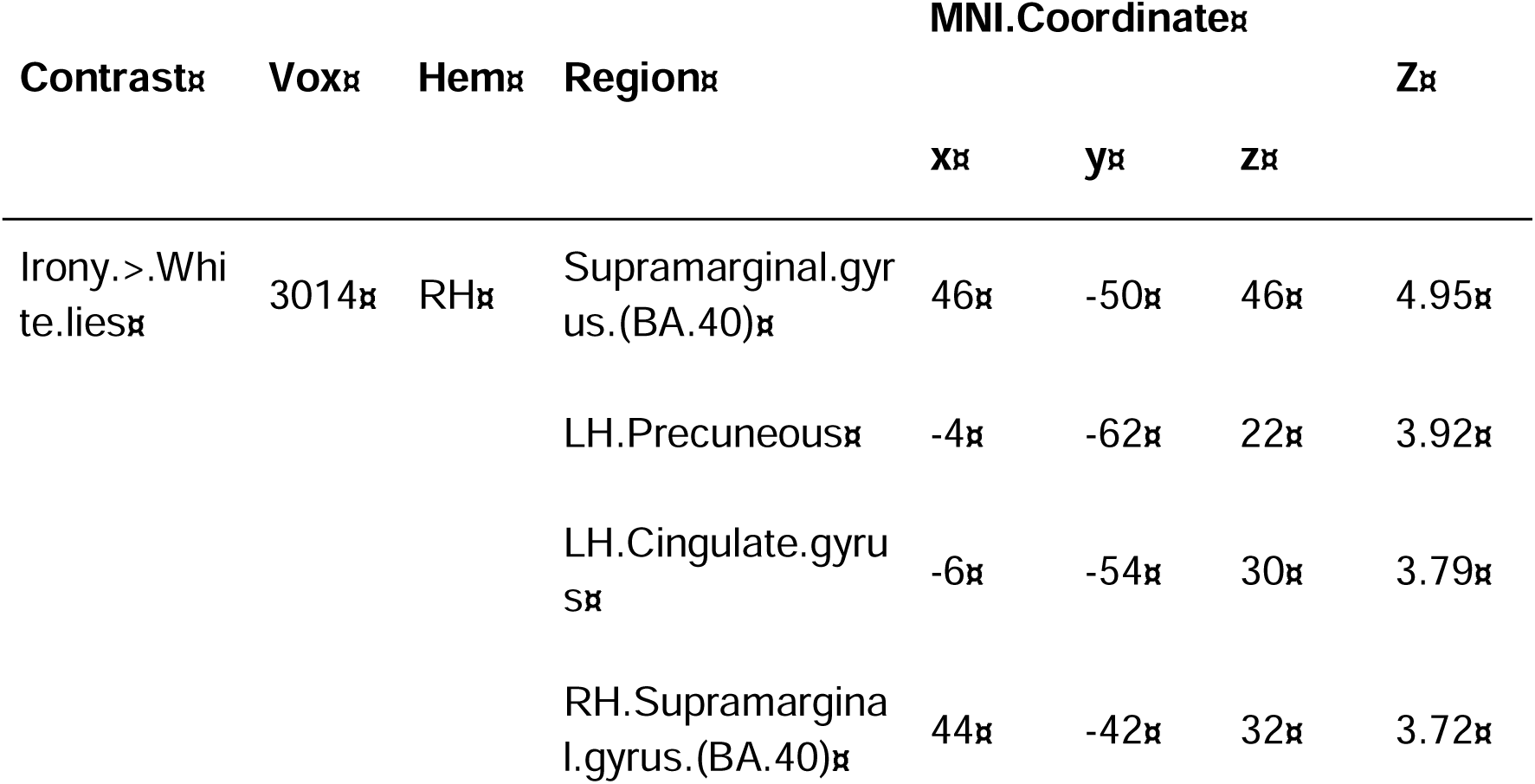
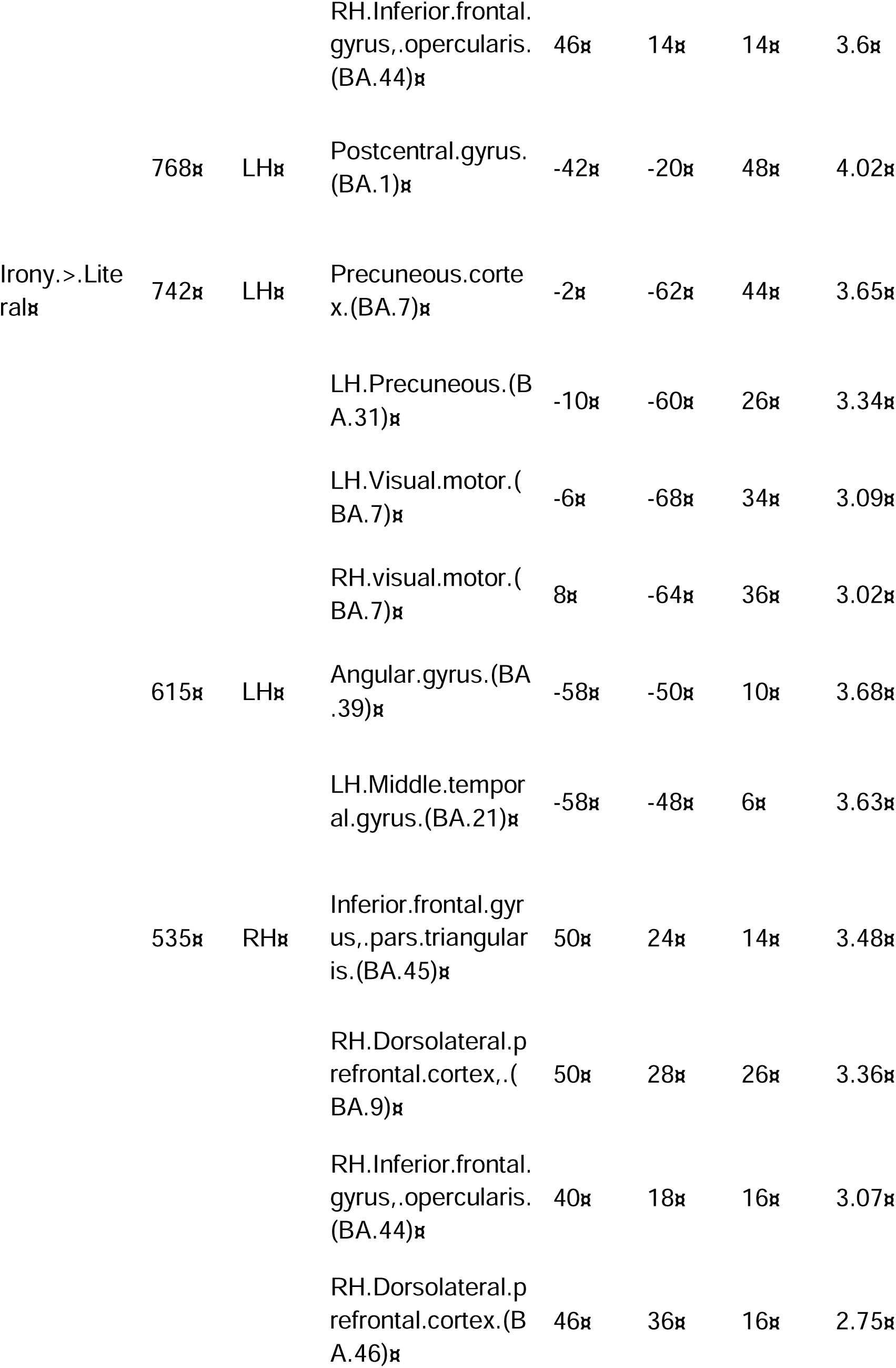
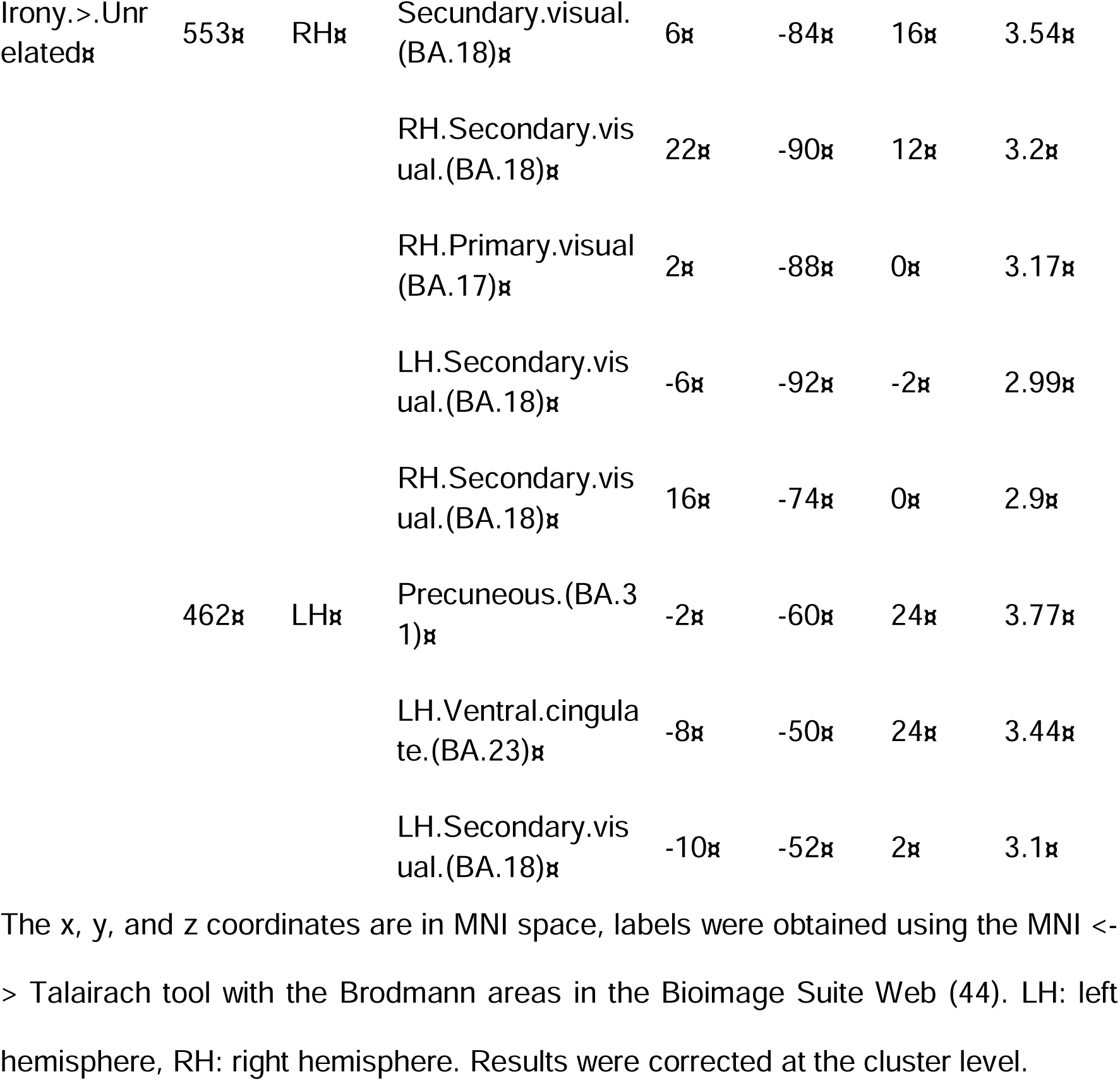
Brain areas exhibiting significant activation in whole brain analysis for contrasts between ironic statements and the other categories, according to GLM analysis. The x, y, and z coordinates are in MNI space, labels were obtained using the MNI <-> Talairach tool with the Brodmann areas in the Bioimage Suite Web (44). LH: left hemisphere, RH: right hemisphere. Results were corrected at the cluster level.

The irony > unrelated contrast showed significant differential activation in two clusters, the right cuneal cortex (BA 18), extending into the associative visual area (BA 18) and lingual gyrus (BA 17) and secondary visual cortex (BA 18); the precuneus (BA 31), extending into the ventral cingulate gyrus (BA 23) and the lingual gyrus (BA 18) (see Fig 5B; Table 3 for MNI coordinates).

The irony > white lies contrast showed significant differential activation in two clusters, the right supramarginal gyrus (SMG; BA 40), extending into the pSTG (BA 22), precuneus (BA 31), cingulate gyrus (BA 31), and the IFG (pars opercularis, BA 44); and the left primary sensory cortex (BA 1,), extending into BA 5 of the primary association cortex. (see Fig 5C; Table 3 for MNI coordinates).

#### Percentage of BOLD signal and cognitive processes

As described in the methods, to analyze the relationship between the activation in the brain during the presentation of the statements and the cognitive processes we measured, the percentage change in the BOLD signal was extracted from regions previously associated with irony. The selected ROIs were: the dorsomedial PFC, IFG, pSTG, and rostromedial PFC, and the signal was extracted from the contrast that compared each statement category (ironic, white lies, literal and unrelated) with the fixation cross.

Scores on social cognition tests predicted BOLD signal change in the IFG during literal (F(2,41) = 6.14, p < 0.01, adjusted-R2 = 0.19), unrelated (F(2,41) = 4.13, p < 0.05, adjusted-R2 = 0.07) and white lies (F(2,41) = 9.14, p < 0.01, adjusted-R2 = 0.27) detection. For the detection of literal statements, the predictors were RMET scores (t = 2.04, p < 0.05) and RMET rt (t = 3.22, p < 0.01), for unrelated statements, RMET rt (t =2.03, p < 0.05), and for white lies, IRI (t = 1.67, p = 0.10) and RMET rt (t = 3.95, p < 0.001) (Table 4, and S3 Table).

**Table 4.** Stepwise regressors for the percentage of BOLD signals associated with the detection of statement categories in Regions of Interest by cognitive process.

| Model.<br>dependent.variable | Social.cognit<br>ion | Executive.fu<br>nctions | Processing.s<br>tyle | Verbal.abiliti<br>es |
| --- | --- | --- | --- | --- |
| Irony |  |  |  |  |
| LH-dmPFC | 0.088 | 0.038 | 0.036 | n.s. |
| LH-rmPFC | 0.086 | 0.067 | 0.053 | n.s. |
| LH-STG | 0.037 | n.s. | n.s. | n.s. |
| LH-IFG | n.s. | 0.037 | 0.202** | n.s. |
| Literal |  |  |  |  |
| LH-dmPFC | n.s. | 0.031 | n.s. | 0.041 |
| LH-rmPFC | n.s. | 0.043 | n.s. | 0.042 |
| LH-STG | n.s. | 0.064 | n.s. | n.s. |
| LH-IFG | 0.193** | 0.035 | 0.328*** | n.s. |
| Unrelated |  |  |  |  |
| LH-dmPFC | n.s. | 0.073* | n.s. | n.s. |
| LH-rmPFC | n.s. | 0.06 | n.s. | n.s. |
| LH-STG | n.s. | 0.067* | n.s. | n.s. |
| LH-IFG | 0.068* | n.s. | 0.171** | 0.023 |
| White.lies |  |  |  |  |
| LH-dmPFC | n.s. | n.s. | n.s. | n.s. |
| LH-rmPFC | 0.073 | n.s. | n.s. | n.s. |
| LH-STG | n.s. | 0.058 | n.s. | n.s. |
| LH-IFG | 0.275*** | 0.032 | 0.25** | n.s. |

The adjusted-R2 is presented for each model. $p = * < 0.05$ , $** < 0.01$ , $*** < 0.001$ .
n.s. = no significant predictors survived in the stepwise regression

Executive functions predicted BOLD signal in the dmPFC during unrelated (F(1,42) =4.37, p < 0.05, adjusted-R2 = 0.073) and STG detection (F(1,42) = 4.09, p < 0.05, adjusted-R2 = 0.067). The significant predictor for dmPFC (t = 2.09, p < 0.05) and STG (t = 2.02, p < 0.05) was go-no-go.

Perceptual processing also predicted BOLD signal change in the IFG during the detection of ironic (F(2,41) = 6.45, p < 0.01, adjusted-R2 = 0.20), literal (F(3,40) = 7.99, p < 0.001, adjusted-R2 = 0.32), unrelated F(2,41) = 5.43, p < 0.01, adjusted-R2 = 0.17), and white lies F(3,40) = 5.94, p < 0.01, adjusted-R2 = 0.25) statements. The predictors for the detection of ironic statements were the local-global rt difference (t = 3.42, p < 0.01) and the no-switch accuracy (t = −3.09, p < 0.01). The predictors for the detection of literal statements were the PRI (t = 2.26, p < 0.05), local-global mean rt difference (t = 4.5, p <0.001), and no switching accuracy (t = −3.76, p < 0.001). The predictors for the detection of unrelated statements were local-global mean rt difference (t = 3.2, p <0.01) and no switching accuracy (t = −2.3, p < 0.05). The predictors for the detection of white lies were the PRI (t = 2.40, p < 0.05), local-global mean reaction time difference (t = 3.61, p <0.001), and no switching accuracy (t = −3.30, p < 0.01). No other test scores were significant predictors of BOLD signal change in other ROIs.

## Discussion

The aims of this paper were to evaluate the cognitive processes and the neural areas associated with the detection of ironic statements. The contextual discrepancy task that we employed showed that ironic statements were difficult to detect, having the lowest accuracy and highest latency of classification (latency). This result agrees with our previous behavioral study (26) and with the proposal that irony is one of the most difficult communication forms (5). With regard to the neural regions involved, we found that the precuneus, which has been associated with ToM, presented consistent activation during the detection of the ironic statements. This activation was shown in the contrast of irony compared to all the other statements, the contrast of irony compared to literal statements, and to unrelated statements. These results are consistent with those of Shibata et al. (2010) and Varga et al. (2013), who studied irony comprehension in Japanese, and Hungarian participants, respectively.

To discover which cognitive processes could be involved in detecting irony, we used a battery of tests that evaluated the domains of social cognition, executive functions, perceptual processing, and verbal abilities. These processes have been previously proposed as pertinent (2,8) and found to be relevant in experimental studies in clinical and non-clinical populations (6,14,15). In the social cognition domain, as was expected, ToM predicted the accuracy and latency of the classification of the ironic statements. The other statement categories, unrelated and white lies, were also predicted by ToM; this relation was not seen for the literal statements. Perhaps ToM facilitates the detection of nonliteral statements, especially because their intended meaning is not immediately apparent since it varies according to the context (8,10,46–48). Interestingly, the AQ questionnaire predicted the nonliteral statements’ classification latency. This questionnaire evaluates abilities associated with autism, such as the degree of social abilities, communication, attention to detail, attention switching, and imagination (17). We found that participants with better abilities, as evaluated by the AQ, took less time to detect nonliteral statements. Together these results show that social cognitive processes play a fundamental role in understanding linguistic information according to the context.

Regarding executive functions, inhibitory control predicted the accuracy and latency of the classification of white lies. The fact that white lies are rated as less sincere than other statements (26) suggests that inhibition of a preponderant interpretation plays an important role (49).

Concerning verbal abilities, the VCI was a positive predictor for detecting ironic, unrelated, and white lies statements. These non-literal statements bear indirect messages and thus require substantial linguistic abilities (7). Indeed, neuroimaging studies have shown that nonliteral language recruits areas associated with language and their homologous such as the right IFG (46).

As for the results of the whole-brain neuroimaging analysis, the contrast between irony and all other statements, as well as between irony and each statement category, showed significant differential activation in the left precuneus, a region that has been associated with ToM, and is a key region of the default mode network that “simultaneously interacts with both the default-mode and frontoparietal networks to distinguish distinct cognitive states” (50). The precuneus seems to be involved in mental imagery and the construction of inferential meanings (9,51–56). This brain region also showed differential activation during speech act recognition, consistent with its role in inferential processes (57). It is worth noting that the precuneus did not show differential activation when detecting other statement categories. Instead, the contrast of the literal and unrelated statements versus the other categories showed differential activation in areas associated with motor activation (e.g., BA 4 and 6). The contrast of white lies versus all other statement categories did not show a differential activation pattern.

We expected that the detection of irony would show significant activation in those areas found by a previous meta-analysis (24), i.e., the MPFC, dorsomedial, and rostromedial (BA 9); IFG (BA 44), and STG (BA 22). Indeed, we found differential activation in the right IFG in the contrasts between irony versus literal statements and white lies. This region has been associated with processes related to language, such as the detection of unusual or incongruous semantic fields (58,59), detection of semantic violations, reanalysis of meanings (60), the integration of linguistic information into context (61), and the selection of the best interpretation according to the social context (62). Event-related potentials (ERPs) studies have shown an increase in the N400, a potential associated with detecting anomalous linguistic information when evaluating irony based on prosody or context (63–65).

The contrast between ironic and literal statements also showed activation in the pSTG. This area has been associated with processes related to social communication, such as ToM (51,53,54), action observation (66), extracting social information about a situation (62,67) and the integration of paralinguistic cues with contextual information (62). ERPs studies of irony employing a variety of contexts and sentences have shown an increase in the P600, which has been associated with integrating information to facilitate irony detection (64,68,69). While eye-tracking studies using context or stories and statements have reported an increment in regressions during the reading of ironic stimuli, the regression has been associated with the integration of necessary information to infer irony (70–73). Together, these findings underscore the relevance of information integration for irony detection.

In addition, the contrast of ironic versus literal statements showed differential activation in the right dlPFC. This region has been associated with the processes of goal-oriented attention, task switching, planning, novelty seeking (74), problem-solving, and working memory (74,75). This region has also been associated with those aspects of complex language processing, such as ambiguity, nonliteral meaning and when cues need to be integrated to infer the linguistic meaning (76).

Compared to the neuroimaging results obtained in the contrast between irony and the other statement categories, the contrasts between literal and unrelated versus all the other categories (e.g. literal > irony + unrelated + white lies), showed differential activation in the primary motor and premotor areas. The primary motor cortex (BA 4) has been associated with the observation of motor tasks, integration of multiple sensory inputs, the inhibition of involuntary movement, and imagining motor sequences (77). The premotor cortex (BA 6) has been associated with the observation of goal-directed actions (78–80), which allows recognition of actions carried out by others; the mirror neuron system, involved in social interactions and attribution of intentions to others (77,80), and recognition of multimodal actions (78). The differential activation of these regions during the detection of literal and unrelated statements versus the other categories suggests that their detection requires additional resources, such as the inhibition of involuntary movement, imagining motor sequences, or observation of actions.

One of our hypotheses was that a global perceptual processing style–central coherence–that focuses on broad patterns of information would be associated with detecting irony. Indeed, the perceptual processing scores predicted the percent change in the BOLD signal in the IFG during ironic, literal, unrelated, and white lies statements detection; the local-global test was the significant predictor. Although it did not significantly predict behavioral performance on the contextual discrepancy task (i.e., accuracy and latency of classification). This result agrees with previous findings associating the IFG with sentence-context integration (61). Also, it could reflect an essential role of a global perceptual processing style in detecting linguistic information during social interactions.

Taken together, these behavioral and neuroimaging results agree with Pexman’s (2008) constraint satisfaction model in that irony detection is predicted by ToM and verbal abilities. In terms of psycholinguistic properties relevant to the detection of irony, Attardo (2000) proposes that ironic statements are relevant — have relation to the context—, are inappropriate to the context, and are meant to convey the intended meaning to the listener. To detect irony, then, a listener needs to compare the statement to the contextual cues present and infer the speaker’s intention. In the contextual discrepancy task, the context varied to give the statement its intended meaning while the statements remained the same. The neuroimaging contrasts between irony and the other statement categories consistently showed the activation of the precuneus, suggesting that this region is involved in comparing context and statement and the inference about the speaker’s intention. It is intriguing that in the contrast of irony versus white lies, but not in the opposite contrast, the SMG showed differential activation. This region appears to be involved in deception (81). These two statement categories are similar in that they give relevant but inappropriate information in relation to the context but differ in terms of the speaker’s intention. Indeed the contrast between white lies and the other categories showed no differential pattern of brain activation, suggesting that the less sincere intention of the speaker, which is the essential element in white lies, was not distinctive enough.

## Conclusion

In conclusion, selecting a heterogeneous sample of participants allowed us to discern the effect of different cognitive processes on irony detection. The behavioral results (accuracy and classification latency) showed that detecting nonliteral statements (irony, unrelated, and white lies) requires ToM and verbal abilities. In addition, white lies detection seemed to involve inhibitory control. Regarding the brain regions associated with irony detection, differential activation patterns were found in areas associated with the ToM (i.e., precuneus) and language (i.e., IFG and pSTG), as was expected based on a previous meta-analysis of pragmatic language (24). Conversely, detecting literal and unrelated statements recruited motor areas (e.g., PMA, PrMA, visual-motor area). No differential activation was found for white lies, possibly because their detection engages processes similar to those for the other categories, with the exception of the deceitful intention. Finally, a global perceptual processing style predicted the percent change in the BOLD signal in the IFG for all the nonliteral and literal statements. For future studies, it would be useful to evaluate the effect of psycholinguistic properties during irony detection. For example, how ironic, sincere, appropriate, or relevant a statement was perceived, and its relation to changes in BOLD signal. Also, including participants with difficulties in understanding social communication, such as ASD, would help to increase the sample variability and improve our understanding of the relation between linguistic, behavioral, cognitive, and brain correlates of irony detection. Finally, our stimuli involve contextual discrepancy for their detection, but during social communication, other cues are present, such as facial expression and prosody. Using other stimuli to present the statements, such as videos, would help to study irony detection in a more ecological manner.

## Data Availability Statement

The data files generated and analyzed in this study are publicly available from the OpenNEURO database (doi:10.18112/openneuro.ds004533.v1.0.0).

## Abbreviations

ToM: Theory of Mind
EF: Executive functions
WCC: weak central coherence
ADHD: attention-deficit hyperactivity disorder
ASD: Autism Spectrum Disorder
AQ: *Autism Spectrum Quotient*
MPFC: medial prefrontal cortex
IFG: inferior frontal gyrus
IPL: inferior parietal lobe
dlPFC: dorsolateral prefrontal cortex
dmPFC: dorsomedial prefrontal cortex
dmPFC: rostromedial prefrontal cortex
SST: Short Story Task
RMET: Reading the Mind in the Eyes Test
IRI: Interpersonal Reactivity Index
SSS: sarcasm self-report scale was used
PEBL: Psychology Experiment Building Language software
PRI: Perceptual Reasoning Index
WAIS-IV: Wechsler Adult Intelligence Scale, fourth edition (WAIS-IV)
VCI: verbal comprehension index
IQR: interquartile range
rt: reaction time
FSL: FMRI Software Library
GLM: general linear model
ROIs: Regions of Interest (ROIs)
BA: Brodmann’s area
MNI: Montreal Neurological Institute
PrMA: premotor area
PMA: primary motor area
VMA: ventral motor area
SMG: supramarginal gyrus
ERPs: Event-related potentials.

## Acknowledgements

We would like to thank all volunteers for their participation in this study. We acknowledge the guidance of Luis Concha Loyola and Juan Fernandez Ruiz on the experimental design and data analysis. For technical support, we thank Erick Pasaye, Leopoldo González, and the technical staff at the Lavis (UNAM-Juriquilla). For experimental support, we want to thank Abril Dominguez. For their careful review of the manuscript, we want to thank Jessica González Norris and Mike Jeziorski.

## Author Contributions

**Conceptualization:** Elizabeth Valles-Capetillo, Magda Giordano

**Data curation:** Elizabeth Valles-Capetillo

**Formal analysis:** Elizabeth Valles-Capetillo, Magda Giordano

**Funding acquisition:** Magda Giordano

**Investigation:** Elizabeth Valles-Capetillo, Cristian Ibarra

**Methodology:** Elizabeth Valles-Capetillo, Magda Giordano

**Project administration:** Elizabeth Valles-Capetillo, Magda Giordano

**Resources:** Magda Giordano

**Supervision:** Elizabeth Valles-Capetillo, Magda Giordano

**Validation:** Elizabeth Valles-Capetillo

**Visualization:** Elizabeth Valles-Capetillo

**Writing – original draft:** Elizabeth Valles-Capetillo, Magda Giordano, Cristian Ibarra

**Writing – review & editing:** Elizabeth Valles-Capetillo, Magda Giordano

