## Supplemental Table 1 for "Irony detection engages the Precuneus and Inferior Frontal Gyrus and relies on integration of contextual cues and inferential skills"

| task | n | min | max | median | iqr | mean | sd | se | ci |
|---|---|---|---|---|---|---|---|---|---|
| SST | 45 | 2.00 | 38.00 | 16.00 | 8.00 | 15.356 | 7.637 | 1.138 | 2.294 |
| SST-SI | 45 | 3.00 | 18.00 | 8.00 | 4.00 | 7.978 | 2.743 | 0.409 | 0.824 |
| SST-COMP | 45 | 277.00 | 336.00 | 311.00 | 10.00 | 309.600 | 10.332 | 1.540 | 3.104 |
| SST-MSR | 45 | 15.00 | 92.00 | 41.00 | 13.00 | 40.200 | 13.829 | 2.062 | 4.155 |
| RMET | 45 | 0.32 | 1.00 | 1.00 | 0.03 | 0.948 | 0.141 | 0.021 | 0.042 |
| RMET-RT | 45 | 0.38 | 0.98 | 0.95 | 0.05 | 0.909 | 0.136 | 0.020 | 0.041 |
| IRI | 45 | -3.29 | 2.81 | 0.93 | 0.40 | 0.818 | 0.786 | 0.117 | 0.236 |
| SSS | 45 | 12.00 | 218.00 | 173.00 | 14.00 | 168.956 | 31.487 | 4.694 | 9.460 |
| AQ | 45 | 66.00 | 120.00 | 107.00 | 11.00 | 104.889 | 10.021 | 1.494 | 3.011 |
| Tower of London | 45 | 9.00 | 32.00 | 27.00 | 5.00 | 25.511 | 4.975 | 0.742 | 1.495 |
| gonogo | 45 | 163.00 | 624.00 | 307.00 | 112.00 | 314.400 | 94.995 | 14.161 | 28.540 |
| Digit span | 45 | 1.12 | 5.38 | 3.12 | 1.69 | 3.266 | 1.090 | 0.162 | 0.327 |
| nback | 45 | 8.00 | 26.00 | 16.00 | 5.00 | 17.067 | 3.922 | 0.585 | 1.178 |
| WAIS-VCI | 45 | 2.00 | 10.00 | 8.00 | 2.00 | 8.467 | 1.502 | 0.224 | 0.451 |
| WAIS-PRI | 45 | 2.00 | 15.00 | 8.00 | 4.00 | 8.133 | 2.989 | 0.446 | 0.898 |
| LG.acc.switch | 45 | 0.00 | 2.00 | 0.00 | 1.00 | 0.467 | 0.548 | 0.082 | 0.165 |
| LG.acc.noswitch | 45 | 2.00 | 12.00 | 9.00 | 2.00 | 8.800 | 2.455 | 0.366 | 0.738 |
| LG.mean.rt.diff | 45 | 85.00 | 129.00 | 102.00 | 12.00 | 103.556 | 11.234 | 1.675 | 3.375 |
